# Eugenol induces potent vasorelaxation in uterine arteries from pregnant goats – A promising natural therapeutic agent for hypertensive disorders of pregnancy

**DOI:** 10.1101/2020.09.22.307629

**Authors:** SC Parija, H Jandhyam, BP Mohanty, P Parasar, NR Nayak

**Author notes:** Corresponding Author: Email address (SCP). These authors contributed equally to this work (SC Parija, H Jandhyam). These authors also contributed equally to this work (BP Mohanty, P Parasar, NR Nayak).

## Abstract

Hypertensive disorders of pregnancy, including preeclampsia, affect about 8-13% of pregnancies and are the leading causes of pregnancy related maternal mortality worldwide. Poorly controlled high blood pressure during pregnancy increases the risk of pregnancy complications and development of future cardiovascular diseases. However, the choice for antihypertensive therapy during pregnancy has been limited due to side effects of many commonly used antihypertensive drugs and lack of other proven safe treatment options. Eugenol is a natural phenolic compound and the main component of clove oil. It is known for its antioxidant, anti-inflammatory and vasorelaxant actions. These beneficial effects of eugenol make it as an excellent therapeutic candidate for treatment of hypertensive disorders of pregnancy. Thus, as a first step, we compared the vasorelaxant effect of eugenol on the middle uterine arterial (MUA) rings from pregnant and nonpregnant goats. Additionally, we examined the potential involvement of the transient receptor potential channel 1 (TRPV1) in mediating the actions of eugenol and compared the effects with a known TRPV1 channel agonist, capsaicin. Isometric tension was measured in MUA rings from endometrial-myometrial junctions of pregnant and nonpregnant goats precontracted with phenylephrine, using a highly sensitive isometric force transducer and an automatic organ bath. The concentration-dependent contractile response curves of eugenol were compared to capsaicin, with and without pre-incubation of the MUA rings with a selective and non-selective TRPV1 antagonists, capsazepine (CAPZ) and Ruthenium Red (RR), respectively. Capsaicin induced concentration-dependent vasorelaxation in nonpregnant PE precontracted MUA rings and the concentration-response curve shifted to the right with significantly reduced pIC_50_ and R_max_ values in the presence of CAPZ and RR. The effects were similar in MUA rings from pregnant animals, except that there was a moderate increase in pIC_50_ values in the presence of RR. Similarly, eugenol induced concentration-dependent vasorelaxation in both nonpregnant and pregnant PE precontracted MUA rings and the effects were markedly antagonized by CAPZ and RR. However, compared to capsaicin, the R_max_ of eugenol was increased 31.25% in nonpregnant and 97.99% in pregnant MUA rings. These results suggest that eugenol has highly potent vasorelaxant effect in MUAs and its effect is partly mediated through activation of the TRPV1 channel. Most importantly, its vasorelaxant effect is about three-fold augmented in pregnancy, suggesting its potential value as a nutraceutical agent and therapeutic candidate for treatment of hypertensive disorders of pregnancy.

## Introduction

In pregnancy the hypertensive disorders may be chronic (pre-existing) or gestational. When blood pressure is maintained at 140/95mmHg before pregnancy to 20th week of gestation is designated as chronic. If such blood pressure is prolonged beyond 20th week of pregnancy it is defined as gestational hypertension. The gestational hypertension is resolved within 42 days after delivery [1,2]. Current treatment for hypertensive pregnancy disorders is to control and manage blood pressure and seizures with antihypertensive and magnesium sulphate [3], however, the ultimate option is to deliver foetus and placenta [4]. Blood pressure levels requiring therapy in pregnancy have been set at higher systolic and diastolic levels compared to the general population. Therapeutic intervention is recommended if blood pressure during pregnancy is maintained at ≤ 149/95 mmHg. The first choice of drugs are methyldopa and labetalol followed by nifedipine to treat severe form of hypertension during pregnancy [2, 5]. Other drugs like ACE-inhibitors and angiotensin II receptor blockers (ARBs) is contraindicated during pregnancy [2, 5] due to adverse foetal outcomes like intrauterine growth retardation, neonatal hypotension, oligohydramnios, and patent ductus arteriosus [6, 7]. Similarly, diuretics can cause placental hyper perfusion and β-blockers except labetolol and atenolol has been reported to be associated with intrauterine growth restriction, preterm delivery, neonatal hypoglycemia and bradycardia [1]. Although, plethora of small therapeutic molecules have been developed in treating hypertensive disorders of pregnancy, they have varying success [9-10]. Based on clinical observational studies, practically none of them are evaluated as per the guidelines of rational pharmacotherapy during pregnancy. In recent years, adverse foetal outcomes have resulted from use of non-recommended antihypertensive drugs pregnant women in different countries [11]. In order to overcome such crisis, use of nutraceutical derived antihypertensive agents would be a better option without any risk for adverse foetal outcomes.

Hypertensive disorders of pregnancy include range of conditions which, if not treated, may lead to preeclampsia [12] and eclampsia, a severe form of hypertension and convulsions and may also consequent into coma and death causing an estimated 14% pregnancy-related maternal deaths. Preeclamptic mothers have higher risk of cardiovascular diseases including chronic heart failure, hypertension, and stroke [8]. Preeclampsia may also associate with intrauterine growth restriction (IUGR) which could cause premature births. These pregnancy complications are particularly of importance in developing countries where incidence of preeclampsia is greater and maternal mortality and preterm births are higher compared to that of developed countries [13-15].

While there is no definitive mechanism or aetiology of preeclampsia, common risk factors are pre-existing diabetes [16], obesity [17], African American ethnicity, and family history of preeclampsia [18-19]. During embryo implantation, placental trophoblast cells invade the uterine arteries (spiral arteries in humans) and induce its remodelling, while obliterating the tunica media of the myometrial spiral arteries which allows the arteries to accommodate increased blood flow to nourish the developing foetus [20]. This abnormal spiral artery remodelling was seen and described over five decades ago in pregnant women who were hypertensive [21]. It has since been shown to be the central pathogenic factor in pregnancies complicated by intrauterine growth restriction, gestational hypertension, and preeclampsia [20]. Defective trophoblastic invasion with associated uteroplacental hypoperfusion may lead to preeclampsia and has been supported by animal and human studies [22-23]. Therapeutic vasorelaxation of spiral arteries could improve placental blood flow, reducing risks to the fetus, and potentially alleviate maternal hypertension.

Vascular tone is regulated in part by transient receptor potential Ca^++^ channels, including TRPV1, a polymodal nociceptor member of the TRPV subfamily. TRPV1 can be activated by capsaicin, noxious heat, extracellular protons, vanilloids, and voltage-dependent depolarization [20]. It is predominately expressed in sensory nerves and non-neuronal tissues and can cause vasodilation through calcitonin gene related peptide (CGRP) and mediate vascular functions via baroreceptor reflex [24]. Its functions in various organs include pain perception, attenuation of inflammatory pain, cell migration, and modulation of receptor activity [25]. TRPV1 in the vascular system plays a vital role in regulation of vasoconstriction and vasodilation [26].

Capsaicin (8-methyl-N-vanillyl-trans-6-nonenamide) is the principal active constituent of hot chilli peppers and has been used as a spice for enhancing flavour and as an analgesic [27]. Capsaicin has significant stimulating effects, including stimulation of thermogenesis, catecholamine secretion from the adrenal medulla, adipogenesis via enhancement of energy metabolism, and weight reduction [28]. Capsaicin is also a selective agonist for the TRPV1 cation channel [29] and capsaicin-induced vasorelaxation has been reported in human coronary artery, rat aorta [30], guinea pig ileum [31], human and porcine coronary arteries [32], rat mesenteric artery [33], and goat mesenteric artery [34].

Another naturally occurring compound with vasorelaxant effects is eugenol (4-allyl-2-methoxyphenol), a chief component of clove oil. Its myorelaxant [35] and vasorelaxant effects in rat [36-37], rabbit thoracic aorta, and rat mesenteric vascular bed [38] suggest it could be used in the treatment of hypertension. Due to the presence of a vanillyl moiety like that in capsaicin, eugenol may also cause vasorelaxation via activation of TRPV1 channels [39]. Eugenol may dilate resistant arteries by increasing the expression of its target genes, sarco/endoplasmic reticulum Ca^2+^-ATPase (SERCA) and big potassium (BK) calcium activated potassium channels (KCa 1.1 or KCNMA1 gene) channel [40-41], thereby relaxing vascular smooth muscle cells and decreasing blood pressure. Information regarding the effects of capsaicin and eugenol on uterine arteries via activation of TRPV1 channels is lacking. Utilizing a novel *Capra hircus* MUA denuded from endo-myometrial junction, a novel arterial model similar to human uterine arteries, and combining functional and electrophysiological studies, we demonstrate that naturally occurring compounds, capsaicin and eugenol, induce vasorelaxation in pregnant and nonpregnant goat MUA and this effect is mediated through endometrial TRPV1. Especially, eugenol due to its higher effectiveness could be used therapeutically to control hypertension and is a potential candidate to treat effects of preeclampsia associated with reduce spiral artery blood flow.

## Materials and methods

### Ethical guidelines

Protocols were approved by the institutional ethical committee on Animal Research (433/CPCSEA/CVS vide ID.No.1586(6)/CVS/dt.03.05.2016).

### Drugs and Chemicals

Commercially available phenylephrine (PE), capsaicin, eugenol, and CAPZ were obtained from Sigma-Aldrich, USA, and RR from MP Biochemicals, India. PE was prepared in triple distilled water whereas Capsaicin and CAPZ, were prepared by dissolving in 70% ethanol.

### Preparation of endo-myometrial junction middle uterine artery rings

Non-pregnant and pregnant uteri intact with broad ligaments and uterine arteries were obtained and placed into aerated ice-cold Modified Krebs-Henseleit Saline (MKHS) (118 mM NaCl, 4.7 mM KCl, 2.5 mM CaCl2, 1.2 mM MgSO4, 11.9 mM NaHCO3, 1.2 mM KH2PO4, and 11.1 mM dextrose, pH 7.4). The arteries from myometrial and endometrial junction were promptly excised. Secondary branches of uterine arteries supplying the uterine horns were carefully cleansed off of fascia and connective tissue in MKHS solution under continuous aeration. The arteries were cut into ∼2 mm long segments of circular rings.

### Isometric tension assay

Endothelium intact middle uterine arterial rings were mounted between two fine stainless-steel L-shaped hooks [42] and kept under a resting tension of 1.5 gm at 37.0±0.5 °C in a thermostatically controlled automatic organ bath (Pan Lab) of 20 mL capacity aerated with carbogen (95% O_2_, 5% CO_2_). Change in isometric tension was measured by a highly sensitive isometric force transducer (Model: MLT0201, AD instrument, Australia) and analysed using Chart 7.1.3 software.

### Experimental protocols

The arterial rings were precontracted by exposure to PE (10^−5^M) to produce a maximal, sustained contraction, then challenged with either cumulative concentrations of Capsaicin (10^−9^ to 10 x 10^−5^ M) or Eugenol (10^−9^ to 10^−5^ M). To investigate the involvement of TRP channels in Eugenol and Capsaicin responses, arterial rings were pre-treated with CAPZ (10^−6^ M), specific blocker of TRPV1, or RR (10^−6^ M), a non-selective TRP channels blocker, for a period of 30 minutes. The concentration-response curves (CRCs) for capsaicin- and eugenol-treated arterial rings were compared with those of untreated controls. The CRCs for capsaicin and eugenol in the presence and absence of RR and CAPZ were also determined. CRCs were plotted and the pIC_50_ calculated for each condition. The pIC_50_ and R_max_/R_Bmax_ of the CRCs were compared for each set of experiments.

### Data analysis and statistics

Data are presented as the mean ± S.E.M. Concentration-response curves are based on the percent relaxation from the agonist-induced contraction (control), and the curves were fitted using interactive non-linear regression in Graph Pad Prism (Graph Pad Prism Software, San Diego, CA, USA). The R_max_/R_Bmax_, mean threshold concentration, and −logIC_50_/pIC_50_ were calculated in Graph Pad Prism. Unpaired Student’s t-test was used to compare groups. A value of p < 0.05 were considered statistically significant.

## Results

### Capsaicin-induced vasorelaxation of middle uterine arteries

Middle uterine arterial rings from endometrial and myometrial junction were mounted and a contraction was induced with PE. Bath application of capsaicin caused concentration-dependent relaxation of this contraction. Capsaicin caused concentration-related vasorelaxation in PE pre-contracted MUA rings of nonpregnant animals (R_max_ = 28.06±0.67%, pIC_50_ = 7.28±0.06; n = 6) (Table 1). When MUA were pre-incubated for 30 min with CAPZ (10^−6^ M), it inhibited Capsaicin-induced relaxation as demonstrated by the rightward shift in the CRC and statistically significant decrease in R_Bmax_ and pIC_50_ (7.52±0.2% and 5.68±0.04, respectively; P<0.001) (Fig 1A). Similarly, pre-incubation with RR shifted the CRC to the right with significant decreases in R_Bmax_ (4.71±0.11%; P<0.001) and pIC_50_ (6.88±0.12; P<0.05) (Fig 1A).

**Table-1.**
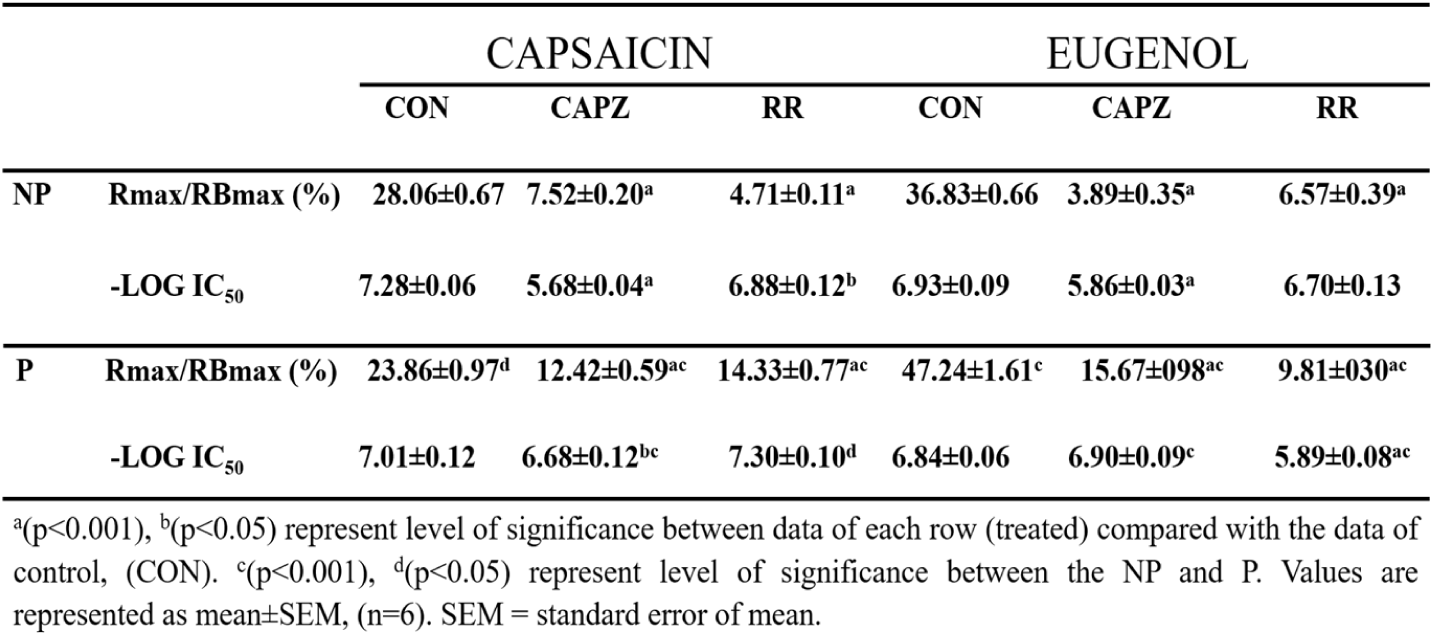
Capsaicin- and eugenol-induced vasorelaxation in phenylephrine (10^−5^M)-precontracted MUA rings from nonpregnant (NP) and pregnant (P) *Capra hircus* in the absence (control) or presence of capsazepine (CAPZ) (10^−6^M) or ruthenium red (RR) (10^−6^M).

|  |  | CAPSAICIN |  |  | EUGENOL |  |  |
| --- | --- | --- | --- | --- | --- | --- | --- |
|  |  | CON | CAPZ | RR | CON | CAPZ | RR |
| NP | Rmax/RBmax (%) | 28.06±0.67 | 7.52±0.20<sup>a</sup> | 4.71±0.11<sup>a</sup> | 36.83±0.66 | 3.89±0.35<sup>a</sup> | 6.57±0.39<sup>a</sup> |
|  | -LOG IC<sub>50</sub> | 7.28±0.06 | 5.68±0.04<sup>a</sup> | 6.88±0.12<sup>b</sup> | 6.93±0.09 | 5.86±0.03<sup>a</sup> | 6.70±0.13 |
| P | Rmax/RBmax (%) | 23.86±0.97<sup>d</sup> | 12.42±0.59<sup>ac</sup> | 14.33±0.77<sup>ac</sup> | 47.24±1.61<sup>c</sup> | 15.67±0.98<sup>ac</sup> | 9.81±0.30<sup>ac</sup> |
|  | -LOG IC<sub>50</sub> | 7.01±0.12 | 6.68±0.12<sup>bc</sup> | 7.30±0.10<sup>d</sup> | 6.84±0.06 | 6.90±0.09<sup>c</sup> | 5.89±0.08<sup>ac</sup> |

<sup>a</sup>( $p < 0.001$ ), <sup>b</sup>( $p < 0.05$ ) represent level of significance between data of each row (treated) compared with the data of control, (CON). <sup>c</sup>( $p < 0.001$ ), <sup>d</sup>( $p < 0.05$ ) represent level of significance between the NP and P. Values are represented as mean±SEM, ( $n=6$ ). SEM = standard error of mean.

**Figure 1.**
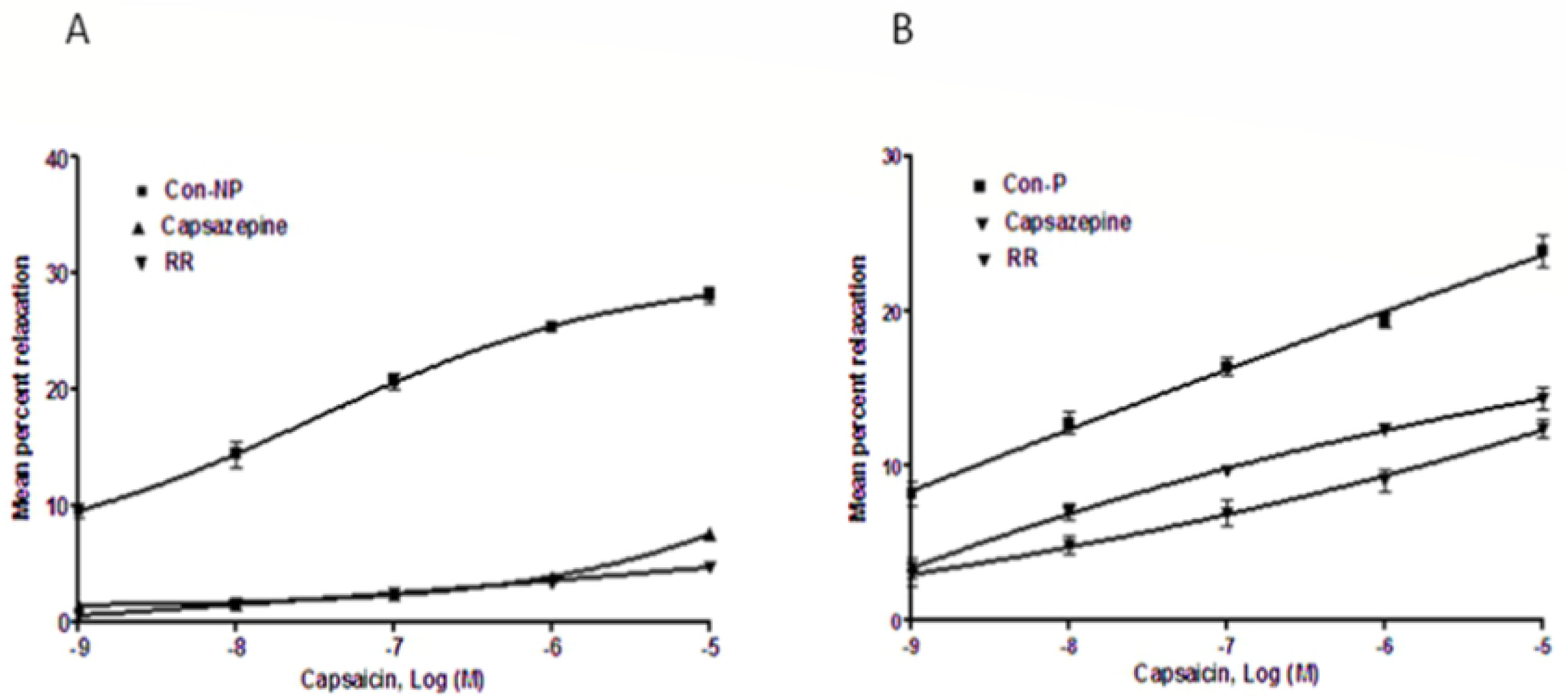

In pregnant goats, capsaicin induced concentration-dependent relaxation in the arterial rings (R_max_ = 23.86± 0.97%, pIC_50_ = 7.01±0.12) (Table 1). When preincubated with CPZ, capsaicin-induced relaxation reduced with a rightward shift in the curve and significant decrease in R_Bmax_ and pIC_50_ (12.42±0.59% and 6.68±0.12, respectively, <0.001) (Fig 1B). In the presence of RR, capsaicin-induced relaxation decreased with a shift of the curve to the right and a significant decrease in R_Bmax_ (14.33±0.77%; P<0.001) and a non-significant increase in pIC_50_ (7.30±0.10) in the presence of RR (Fig 1B).

### Vasorelaxation induced by Eugenol involves TRPV1 channel activation

MUA rings of nonpregnant uteri when incubated with eugenol significantly relaxed the arterial rings precontracted with PE (R_max_ = 36.83±0.66%, pIC50 = 6.93±0.09) (Table 1). To investigate the endothelial-cell dependent mechanism of Eugenol-induced vasorelaxation, the experiments were performed in the presence of TRPV1 inhibitors CAPZ and RR. The eugenol CRC shifted to the right with a significant decrease in R_Bmax_ and pIC_50_ in the presence of CAPZ (R_Bmax_ = 3.89±0.35%, pIC_50_ = 5.86±0.03; P<0.001; n=6) (Fig 2A). On the other hand, incubation with RR shifted curve rightward with significant decrease in R_Bmax_ (6.57±0.39%; P<0.001) and a nonsignificant decrease in pIC_50_ (6.70±0.13) (Fig2A).

**Figure 2.**
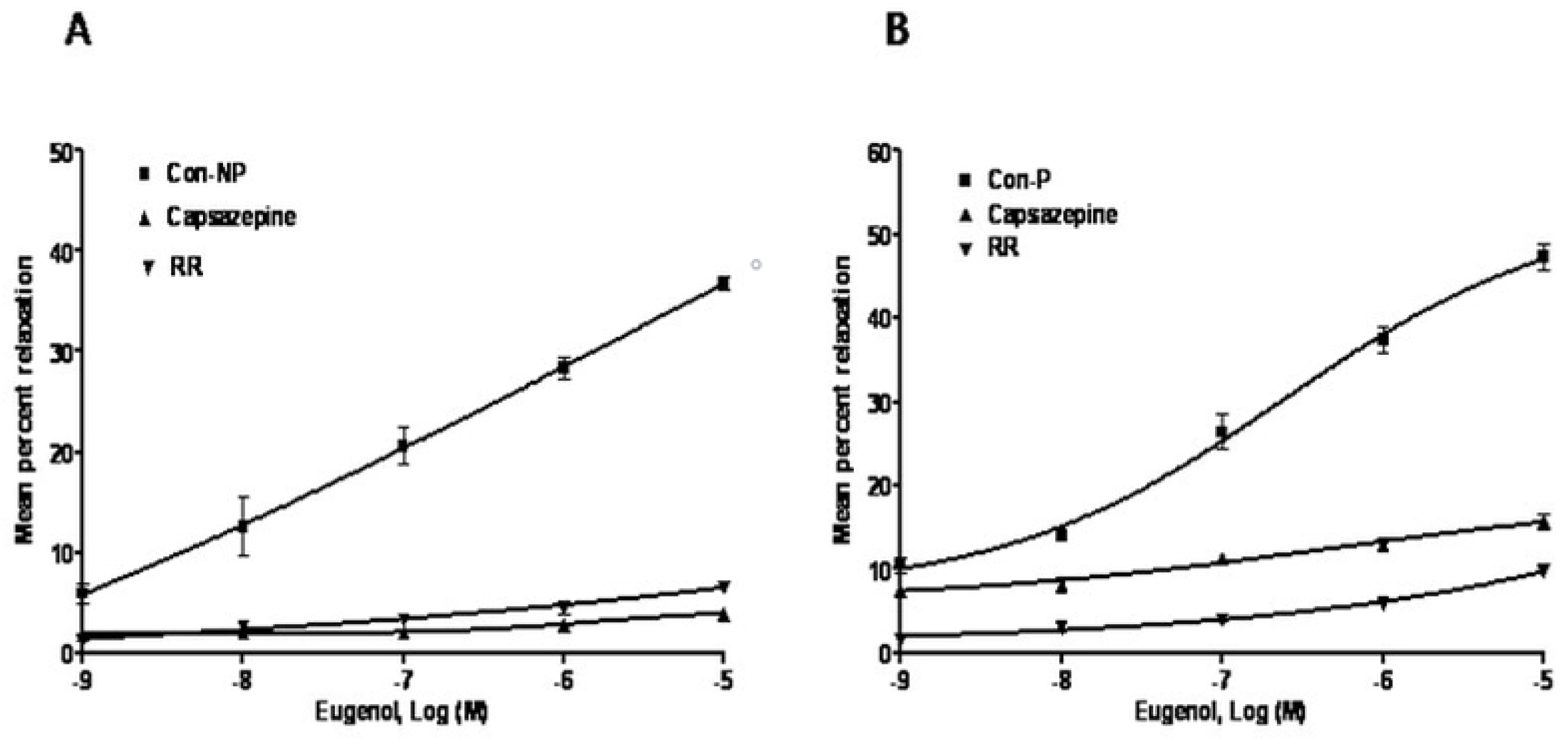

In the MUA of pregnant uteri, eugenol induced vasorelaxation (R_max_ 47.24±1.61%, pIC_50_ 6.84±0.06) (Figure 2B). When incubated with CAPZ, a competitive TRPV1 and TRPM8 receptor antagonist, the curve shifted to the right and caused a significant decrease in R_Bmax_ (15.67±0.98%; P<0.001) and a nonsignificant increase in pIC_50_ (6.9±0.09) (Figure 2B). However, eugenol-induced relaxation significantly decreased (R_Bmax_ 9.81±0.30%, pIC_50_5.89±0.13; P<0.001; n=6) in the presence of RR (Fig 2B). The mean maximal effects of different ligands relative to capsaicin vasorelaxation (100%) are presented in Fig 3.

**Figure 3.**
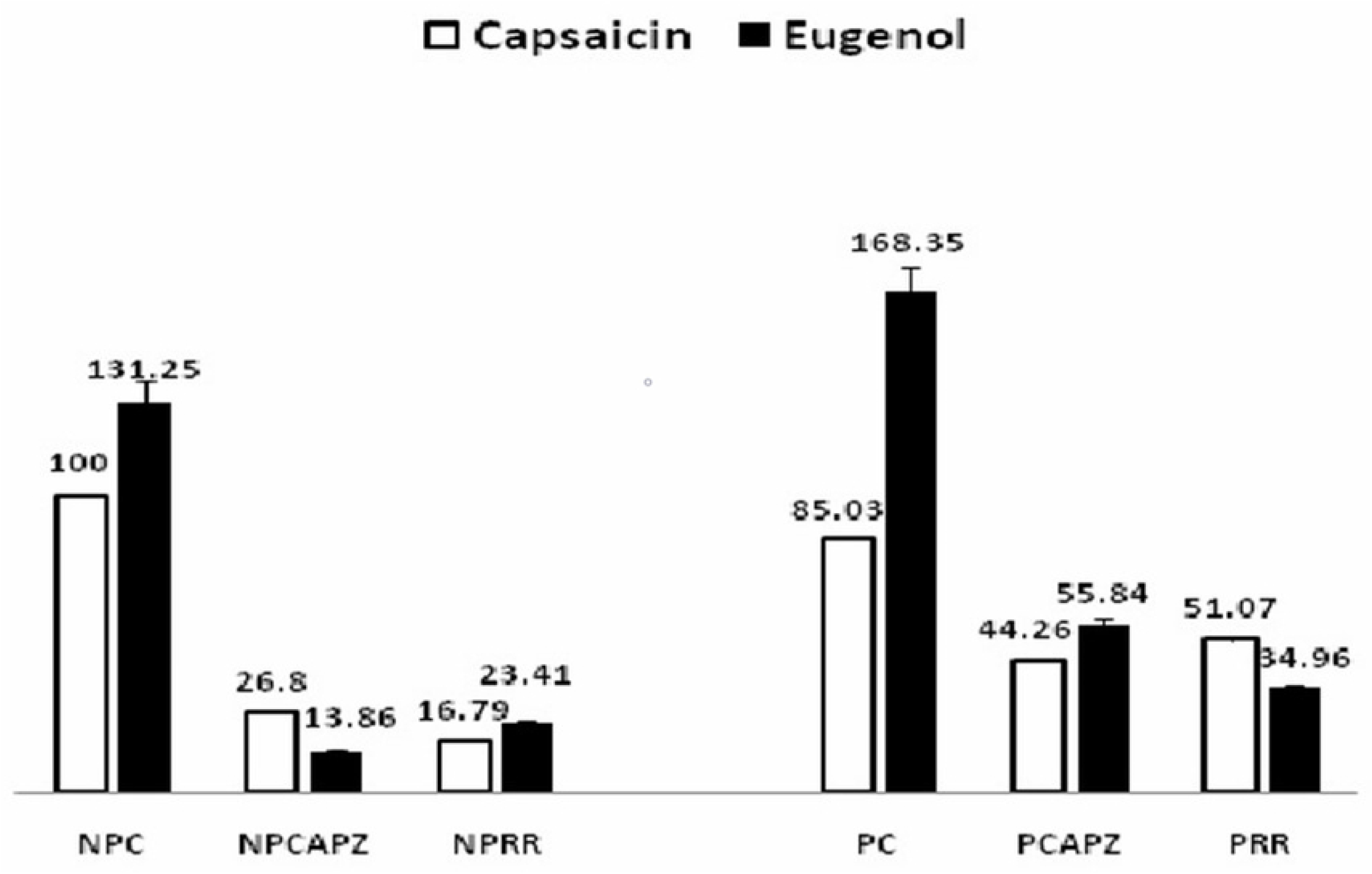

## Discussion

Hypertensive disorders of pregnancy are most common medical disorders of pregnancy which complicate 1 in 10 pregnancies. Notably, preeclampsia, a hypertensive disorder of pregnancy, affects 2–8% of pregnancies worldwide. Whereas the rates of occurrence of preeclampsia overall have remained unchanged, the cases of severe preeclampsia have begun to rise over the last several decades [43-44]. Treatment recommendations for hypertension in pregnancy is set at varying levels than in general population. Blood pressure levels requiring therapy in pregnancy have been set at higher systolic and diastolic levels compared to the general population.

Treatment of severe hypertension requires pharmacological intervention using methyldopa, labetalol, and nifedipine; however, current clinical management is limited, and more innovative medical therapies are needed. Labetalol may be associated with foetal growth restriction and nifedipine may inhibit labour. Hydralazine, other antihypertensive drug, has been tested few controlled trials, and may cause neonatal thrombocytopenia. Beta-receptor blockers may reduce uteroplacental blood flow and have the risk of growth restriction when started in the first and second trimester. It also may cause neonatal hypoglycaemia at higher dose. Hydrocholorothiazide may cause volume contraction and electrolyte disorders, however, may be helpful when combined with methyldopa to mitigate compensatory fluid retention. During pregnancy, the challenge is in deciding when to use antihypertensive medications and what level of BP to target. Despite advancements in therapeutic discoveries, it is crucial to consider the safety of the novel agents as therapeutics in pregnant women and to the foetus. Small molecules, despite their role in clinical management, possess varying success rate. Sildenafil, a phosphodiesterase inhibitor, for instance, was originally thought to be a promising uterine vasodilator. However, it did not improve pregnancy outcomes compared to placebo treatment in the women with intrauterine growth restriction [45]. No antihypertensive drug has been proven safe for use during the first trimester [46].

Other small molecules such as statins (e.g., Pravastatin) because of their stimulatory effects on hemeoxygenase (HO) and improved vascular functions [47] show clinical promise in treating early onset preeclampsia; however, randomized clinical trials are still in progress [45]. Ouabain, a cardiac glycoside, has been shown to inhibit sFLT-1 mRNA and protein expression through the HIF-1α/heat shock protein 27 pathway in human cytotrophoblasts and explant cultures; however, further investigation of ouabain as a therapeutic is yet to be characterized [45-46]. Novel agents acting upon immediate mediators of vasorelaxation such as TRP ion channels are potential therapeutic candidates for preeclampsia and other hypertensive pregnancy disorders.

Therefore, novel natural or endogenous vasorelaxants are needed to treat hypertension during pregnancy with lower risks and better safety during pregnancy. Novel agents acting upon immediate mediators of vasorelaxation such as TRP ion channels are potential therapeutic candidates for preeclampsia and other hypertensive pregnancy disorders. The present study examines the role of natural compounds, capsaicin and eugenol, and assesses the sensitivity of TRPV1 ion channel in our novel model of spiral arteries, the caprine middle uterine arteries. TRP channels, a superfamily of about 28 nonselective cation channels playing key role in vasomotor tone, are divided into 7 subfamilies including TRP Vanilloid (TRPV) [48]. TRPV1 plays key regulatory role in thermal nociception [49] and is important in thermal hyperalgesia after inflammation or tissue injury [50]. TRPV1 channels are found on nociceptive axons in rat vagina [51] and human cervix [52]. In addition, estrogen enhances pain evoked by uterine distension via a TRPV1 receptor-dependent mechanism [53]. Hypertension is the most common medical disorder of pregnancy, and it is possible that activation of the TRPV1 receptor in the uterine vasculature could control such hypertension during pregnancy.

Our study demonstrate that vasorelaxation to capsaicin is attenuated by15% in uterine artery of pregnant as compared to that of non pregnant goat. Eugenol-induced vasorelaxation in PE-precontracted arteries was 31% greater in nonpregnant goats and 68% greater in pregnant goats than that of capsaicin (100%) in nonpregnant goats. These results clearly indicate that the contribution of Capsaicin-sensitive TRPV1 channels to vasorelaxation of uterine artery is reduced in pregnancy.

The TRPV1 antagonist CAPZ is a chemical derivative of capsaicin that has been used extensively as a pharmacological tool to assess the role of TRPV1 in inflammatory hyperalgesia [54]. Similarly, RR, a water soluble polycationic dye, has been found to block the pores of the capsaicin-responsive cation channel TRPV1, thus interfering with all polymodal ways of TRPV1 activation [55]. TRPV1 channel blocker has been reported to inhibit TRPV1-mediated vasorelaxation in human coronary artery at 0.1 mM by 94%, rat skin saphenous nerve preparation at 50 μM, and rat mesenteric artery at 3 μM by about 40% [56]. RR at 10 μM inhibited capsaicin-induced vasorelaxation in goat superior mesenteric artery by 83%, indicating that the TRPV1 channel in this tissue possesses antagonist binding sites with greater sensitivity to RR [34]. TRPV1-induced vasorelaxation may be mediated by activation of endothelial TRPV1 channels, causing increased Ca^2+^ influx and consequent phosphorylation of protein kinase A (PKA) and eNOS, leading to release of the vasorelaxing factors cAMP and nitric oxide [28]. We demonstrate that CAPZ and RR reduced maximal capsaicin-induced uterine artery vasorelaxation by 73% and 83%, respectively, in nonpregnant and by 41% and 34%, respectively, in pregnant goats. The reduced blocking effect of CAPZ and RR at capsaicin sensitive TRPV1 channels in pregnant as compared to that of nonpregnant uterine arteries, RR having a greater inhibitory effect than does CAPZ in nonpregnant goats and CAPZ having a greater inhibitory effect than RR in pregnant goats clearly indicates that there is a significant reduced sensitivity of capsaicin sensitive TRPV1 channels to both the antagonists in uterine artery of pregnant goat. The reduced sensitivity of TRPV1 channels could be resulted from reduced function arising from down-regulation or expression of capsaicin sensitive TRPV1 channels or expression of other members of the TRP families (e.g., TRPM8 and TRPA1) in these remodelled arteries whose roles in female reproductive tract is yet to be determined.

Eugenol has numerous pharmacological effects, including inhibition of voltage-gated sodium channels and activation of TRPV1 [56]. Electrophysiological studies show that eugenol activates Ca^2+^-permeable ion channels [57]. Both the in vitro and in vivo actions of eugenol are reversed by CAPZ, suggesting that its effects are mediated by TRPV1. However, eugenol is a nonselective TRPV1 agonist, as it is also able to activate other TRP channels, such as TRPA1 and TRPM8 [58], and TRPV4 [59]. In this study we observed that eugenol vasorelaxation increased from 31% in nonpregnant goats to 68% in MUA of pregnant caprine when compared to that of capsaicin-induced vasorelaxation (100%). Furthermore, CAPZ and RR inhibited eugenol vasorelaxation (100%) in uterine artery by 90% and 82%, respectively, in nonpregnant and by 67% and 79%, respectively, in pregnant caprine indicating that eugenol vasorelaxation in the MUA of nonpregnant goats is predominantly mediated by capsaicin sensitive TRPV1 channels. This reduced inhibitory effect of CAPZ on eugenol-induced vasorelaxation compared to that of RR clearly suggests that, in pregnant uterine artery, eugenol vasorelaxation could be mediated in part by both CAPZ sensitive channels and CAPZ-resistant TRPV1 channels. Eugenol in MUA of gravid uteri could be activating other TRP channels such as TRPA1 and TRPM8 and this could explain the CAPZ-resistant eugenol vasorelaxation observed in uterine arteries from pregnant animals.

Collectively, our results indicate that capsaicin sensitive TRPV1 channels show reduced sensitivity in the MUA in pregnant goats as compared to those in nonpregnant. We demonstrate that reduced blocking effects and/or binding affinities of both the selective and nonselective TRPV1 antagonists in the MUA in pregnancy could be due to decreased function or expression of capsaicin sensitive TRPV1 during vascular remodelling. Our findings show greater differential increase in vasorelaxation between eugenol and capsaicin in the MUA from pregnant than from nonpregnant animals. Though mechanism is yet to be discovered, eugenol-induced selective vasorelaxation during pregnancy show high impact on the therapeutic potential of this compound. A further investigation to characterize affinity of eugenol to both TRPV1 and other related members of the TRP family in caprine model is needed. Likewise, a greater inhibitory effect by both CAPZ and RR on eugenol-induced vasorelaxation in MUA as compared to that on capsaicin vasorelaxation clearly indicates that the TRPV1 channel has a greater sensitivity to eugenol that is further augmented upon vascular remodelling in pregnancy. TRPV1 shows wide tissue distribution with highest expression in dorsal root ganglion, trigeminal ganglia, and nodose ganglia [20] and is associated with inflammatory conditions and nociception. TRPV1 expression occurs in epidermis [60], polymorphonuclear granulocytes [61], smooth muscles [62], mast cells [63], dendritic cells, and macrophages [64]. Several studies have shown its expression in neuronal tissues brain regions [65, 66], and in non-neuronal tissues Similarly, it is sensitized in the presence of inflammatory mediators such as prostaglandins and bradykinins, physical and chemical stimuli [67-69]. Capsaicin and other related compounds are lipophilic and share structural similarity with endogenous fatty acid derivatives that have been identified as TRPV1 agonists such as anandamide (an endocannabinoid) interact at intracellular regions of TRPV1 [70]. Mild acidic pH at extracellular side of the channel [71] also activates TRPV1. Various activators of TRPV1 potentiate the effect leading to synergistic and enhanced activity [72] which is important in the context of inflammation due to wide variety of inflammatory agents generated in inflamed conditions.

In the context of pregnancy where an inflammatory condition ensues producing cytokines and recruitment of leukocytes to the uterine site for implantation, growth, development, and progression of pregnancy. Hormonal, cytokine, and immune milieu during pregnancy would provide a continuous synergising and activating stimulus to TRPV1 and thus eugenol and capsaicin could impact on this ion channel and induce vasorelaxation during hypertensive pregnancy disorders including preeclampsia. Thus, these compounds can be promising vasorelaxation inducers; however, further mechanistic studies are needed.

## Conclusions

Our results indicate that eugenol and capsaicin can be used as selective vasorelaxant and TRPV1 may be involved in this phenomenon. These results are supported by evidence of induced vasorelxation in the pregnant middle uterine arteries by the agents. We also showed that eugenol is more potent and stronger inducer of vasorelaxation as shown by greater inhibitory effect of CAPZ and RR on eugenol-induced vasorelaxation in pregnant MUA compared to that in nonpregnant animals. Increased sensitivity of TRPV1 to eugenol during pregnancy could be due to increased sensitivity of TRPV1 and/or activation of other TRP channels, such as TRPV4, TRPA1, and TRPM8. Eugenol could be therapeutically useful in treatment and prevention of hypertension during pregnancy (i.e., preeclampsia). Selective vasorelaxation in the pregnant Caprine MUAs in novel spiral artery model by eugenol mediated by TRP channels makes it a potent therapeutic vasorelaxant and begs a further investigation of the underlying mechanisms.

## Acknowledgement

The authors are grateful to the University Grants Commission and Government of India for providing necessary logistical support and a student fellowship grant to JH. We are also thankful to OUAT for providing necessary infrastructure for the smooth conduct of the research.NRN was supported by NICHD grant # R01 HD088549 (the content is solely the responsibility of the authors and does not necessarily represent the official views of the US National Institutes of Health).

## Author Contributions

Conceptualization: Jandhyam H, Parija SC, Nayak NR

Data curation: Jandhyam H, Parija SC

Formal analysis: Jandhyam H

Funding acquisition: Jandhyam H, Parija SC, Nayak NR

Investigation: Jandhyam H, Parija SC

Methodology: Jandhyam H, Parija SC

Project Administration: Parija SC

Resources: Parasar P, Nayak NR, Mohanty BP

Software: Parija SC, Jandhyam H

Supervision: Parija SC, Mohanty BP

Validation: Parija SC, Parasar P, Nayak NR

Writing: Parija SC, Jandhyam H,Parasar P, Nayak NR,Mohanty BP

## References

1. Vest AR, Cho LS. Hypertension in pregnancy. Cardiol Clin. 2012;30(3):407–23.

2. Mancia G, Fagard R, Narkiewicz K, Redón J, Zanchetti A, Böhm M, Christiaens T, Cifkova R, De Backer G, Dominiczak A, Galderisi M, Grobbee DE, Jaarsma T, Kirchhof P, Kjeldsen SE, Laurent S, Manolis AJ, Nilsson PM, Ruilope LM, Schmieder RE, Sirnes PA, Sleight P, Viigimaa M, Waeber B, Zannad F; Task Force Members. 2013 ESH/ESC Guidelines for the management of arterial hypertension: the Task Force for the management of arterial hypertension of the European Society of Hypertension (ESH) and of the European Society of Cardiology (ESC). J Hypertens. 2013;31(7):1281–357. doi: 10.1097/01.hjh.0000431740.32696.cc.

3. Witlin AG, Sibai BM. Magnesium sulfate therapy in preeclampsia and eclampsia. Obstet Gynecol. 1998;92(5):883–9.

4. Lambert G, Brichant JF, Hartstein G, Bonhomme V, Dewandre PY. Preeclampsia: an update. Acta Anaesthesiol Belg. 2014;65(4):137–49.

5. World Health Organization. WHO Guidelines Approved by the Guidelines Review Committee. WHO Recommendations for Prevention and Treatment of Pre-Eclampsia and Eclampsia; 2011. Available from: www.ncbi.nlm.nih.gov/pubmed/23741776.

6. Quan A. Fetopathy associated with exposure to angiotensin converting enzyme inhibitors and angiotensin receptor antagonists. Early Hum Dev. 2006;82(1):23–8.

7. Al-Maawali A, Walfisch A, Koren G. Taking angiotensinconverting enzyme inhibitors during pregnancy: is it safe? Can Fam Physician. 2012;58(1):49–51.

8. Meads CA, Cnossen JS, Meher S, Juarez-Garcia A, ter Riet G, Duley L, Roberts TE, Mol BW, van der Post JA, Leeflang MM, et al. Methods of prediction and prevention of preeclampsia: systematic reviews of accuracy and effectiveness literature with economic modelling. Health Technol Assess. 2008;12(6):iii–v. 1-270.

9. Rolnik DL, Wright D, Poon LC, et al. Aspirin versus placebo in pregnan-cies at high risk for preterm preeclampsia. N Engl J Med. 2017;377:613–22.

10. Caritis S, Sibai B, Hauth J, Lindheimer MD, Klebanoff M, Thom E, VanDorsten P, Landon M, Paul R, Miodovnik M, Meis P, Thurnau G. Low-dose aspirin to prevent preeclampsia in women at high risk. National Institute of Child Health and Human Development Network of Maternal-Fetal Medicine Units. N Engl J Med. 1998;338:701–5.

11. Brown CM, Garovic VD. Drug treatment of hypertension in pregnancy. Drugs. 2014;74(3):283–96. doi: 10.1007/s40265-014-0187-7

12. Al Khaja KA, Sequeira RP, Alkhaja AK, Damanhori AH. Drug treatment of hypertension in pregnancy: a critical review of adult guideline recommendations. J Hypertens. 2014;32(3):454–63.

13. Say L, Chou D, Gemmill A, Tuncalp O, Moller AB, Daniels J, Gulmezoglu AM, Temmerman M, Alkema L. Global causes of maternal death: a WHO systematic analysis. The Lancet. Global health. 2014;2(6):e323–33.

14. McBride CA, Bernstein IM, Badger GJ, Horbar JD, Soll RF. The effect of maternal hypertension on mortality in infants 22, 29weeks gestation. Pregnancy hypertension. 2015; 5(4):362–66.

15. Lain KY, Roberts JM. Contemporary concepts of the pathogenesis and management of preeclampsia. Jama. 2002;287(24):3183–6.

16. Rosenberg TJ, Garbers S, Lipkind H. Chiasson M. A. Maternal obesity and diabetes as risk factors for adverse pregnancy outcomes: Differences among 4 racial/ethnic groups. American J of Public Health. 2005;95(9):1545–51.

17. Boghossian NS, Yeung E, Mendola P, Hinkle SN, Laughon SK, Zhang C, Albert PS. Risk factors differ between recurrent and incident preeclampsia: A hospital-based cohort study. Annals of Epidemiology. 2014;24(12):871–7e3.

18. Lim, K.-H., Steinberg, G., & Ramus, R. M. (2018). Preeclampsia. Retrieved November 14, 2018, from http://emedicine.medscape.com/article/1476919-overview.

19. Duckitt K, Harrington D. Risk factors for pre-eclampsia at antenatal booking: Systematic review of controlled studies. British Medical Journal. 2005;330(7491):565.

20. Kaufmann P, Black S, Huppertz B. Endovascular trophoblast invasion: implications for the pathogenesis of intrauterine growth retardation and preeclampsia. Biol Reprod. 2003;69(1):1–7.

21. Brosens I. A study of the spiral arteries of the decidua basalis in normotensive and hypertensive pregnancies. J Obstet Gynaecol Br Commonw. 1964; 71():222–30.

22. Gilbert JS, Babcock SA, Granger JP. Hypertension produced by reduced uterine perfusion in pregnant rats is associated with increased soluble fms-like tyrosine kinase-1 expression. Hypertension. 2007; 50(6):1142–7.

23. Makris A, Thornton C, Thompson J, Thomson S, Martin R, Ogle R, Waugh R, McKenzie P, Kirwan P, Hennessy A. Uteroplacental ischemia results in proteinuric hypertension and elevated sFLT-1. Kidney Int. 2007 May; 71(10):977–84.

24. Baylie RL, Brayden JE. TRPV channels and vascular function. Acta Physiol (Oxf). 2011;203(1):99–116. doi: 10.1111/j.1748-1716.2010.02217.x

25. Waning J, Vriens J, Owsianik G, Stüwe L, Mally S, Fabian A, Frippiat C, Nilius B, Schwab A. A novel function of capsaicin-sensitive TRPV1 channels: Involvement in cell migration. Cell Calcium. 2007;42:17–25.

26. Sharma SK, Vij AS, Sharma M. Mechanisms and clinical uses of capsaicin. Eur J Pharmacol. 2013;720:55–62.

27. Reyes-Escogido ML, Gonzalez-Mondragon EG, Vazquez-Tzompantzi E. Chemical and pharmacological aspects of capsaicin. Molecules. 2011; 16:1253–70.

28. Yang D, Luo Z, Ma S, Wong WT, Ma L, Zhong J, et al. Activation of TRPV1by dietary capsaicin improves endothelium-dependent vasorelaxation and prevents hypertension. Cell Metab. 2010;12:130–41.

29. Garcia-Sanz N, Fernandez-Carvajal A, Morenilla-Palao C, Planells-Cases R,Fajardo-Sanchez E, Fernandez-Ballester G, Ferrer-Montiel A. Identification of a tetramerization domain in the C terminus of the va-nilloid receptor. J Neurosci. 2004;24:5307–14.

30. Lo YC, Hsiao HC, Wu DC, Lin RJ, Liang JC, Yeh JL, et al. A novel capsaicin derivative VOA induced relaxation in rat mesenteric and aortic arteries: involvement of CGRP, NO, cGMP, and endothelium-dependent activities. J Cardiovasc Pharmacol. 2003;42:511–20.

31. Fujimoto S, Mori M, Tsushima H, Kunimatsu M. Capsaicin-induced, capsazepine-insensitive relaxation of the guinea-pig ileum. Eur J Pharmacol. 2006;530:144–51.

32. Gupta S, Lozano Cuenca J, Villalon CM, de Vries R, Garrelds IM, Avezaat CJ, et al. Pharmacological characterisationof capsaicin-induced relaxations in human and porcineisolated arteries. Naunyn chmiedebergs Arch Pharmacol. 2007;375:29–38.

33. Chen Q, Zhu H, Zhang Y, Zhang Y, Wang L, Zheng L. Vasodilating effect of capsaicin on rat mesenteric artery and its mechanism. J Zhejiang Univ Med Sci. 2013;42:177–83.

34. Mohanty I, Nayak NR, Parija SC. Augmentation of capsaicin-induced vasorelaxation of superior mesenteric artery (Capra hircus) in acidosis: role of TRPV1 channels. Indian Journal of Traditional Knowledge. 2016;15:625–31.

35. Leal-Cardoso JH, Coelho de Souza AN, Souza IT, Figueiredo IM. Effects of eugenol on excitation– contraction coupling in skeletal muscle. Arch. Int. Pharmacodyn Ther. 1994;327:113–24.

36. Leal-Cardoso JH, Lahlou S, Coelho-de-Souza AN, Criddle DN, Pinto Duarte GIB, Santos MIV. Inhibitory actions of eugenol on rats isolated ileum. Can J Physiol Pharmacol. 2002;80:901–6.

37. Damiani CE, Rossoni LV, Vassallo DV. Vasorelaxant effects of eugenol on rat thoracic aorta. Vascul Pharmacol. 2003;40:59–66.

38. Nishijima H, Uchida R, Kameyama K, Ohkubo T, Kitamura K. Mechanisms mediating the vasorelaxing action of eugenol, a pungent oil, on rabbit arterial tissue. Jpn J Pharmacol. 1999;79:327–34.

39. Szallasi A, Cruz F, Geppetti P. TRPV1: a therapeutic target for novel analgesic drugs? Trends Mol Med. 2006;12:545–54.

40. Boominathan. Natural product-based therapy for Hypertension: Eugenol, a natural compound isolated from herb Ocimum sanctum Linn (Tulsi), relaxes vascular smooth muscle cells, and decreases blood pressure by increasing the expression of SERCA and BKCa Channels. Genome-2-Bio-Medicine Discoverycenter. 2015. http://genomediscovery.org.

41. Peixoto-Neves D, Leal-Cardoso JH, Jaggar JH. Eugenoldilates rat cerebral arteries by inhibiting smooth muscle cellvoltage-dependent calcium channels. J Cardiovasc Pharmacol. 2014;64:401–6.

42. Mulvany MJ, Aalkjaer C. Structure and function of small arteries. Physiol Rev. 1990;70:921–961.

43. Phipps E, Prasanna D, Brima W, Jim B. Preeclampsia: Updates in Pathogenesis, Definitions, and Guidelines. Clin J Am Soc Nephrol. 2016;11(6):1102–13. doi: 10.2215/CJN.12081115

44. Ananth CV, Keyes KM, Wapner RJ. Pre-eclampsia rates in the United States, 1980-2010: age-period-cohort analysis. BMJ. 2013;347:f6564.

45. Sharp A, Cornforth C, Jackson R, Harrold J, Turner MA, Kenny LC, Baker PN, Johnstone ED, Khalil A, von Dadelszen P, Papageorghiou AT, Alfirevic Z, STRIDER group. Maternal sildenafil for severe fetal growth restriction (STRIDER): a multicentre, randomised, placebo-controlled, double-blind trial. Lancet Child Adolesc Health. 2018;2:93–102

46. Podymow T, August P. Update on the Use of Antihypertensive Drugs inPregnancy. Hypertension. 2008;51:960–969.

47. Levine RJ, Lam C, Qian C, Yu KF, Maynard SE, Sachs BP, Sibai BM, Epstein FH, Romero R, Thadhani R, Karumanchi SA; CPEP Study Group. Soluble endoglin and other circulating antiangiogenic factors in preeclampsia. N Engl J Med. 2006;355:992–1005.

48. Venkatachalam K., and Montell C, “TRP channels,”AnnualReview of Biochemistry, vol.76, pp.2007; 387–417.

49. Christoph T, Bahrenberg G, De Vry J, Englberger W, Erdmann VA, Frech M, Kogel B, Rohl T, Schiene K, Schroder W, Seibler J, Kurreck J. Investigation of TRPV1 loss-of-function phenotypes in transgenic shRNA expressing and knockout mice. Molecular and Cellular Neuroscience. 2008; 37:579–89. [PubMed: 18249134]

50. Vilceanu D, Honore P, Hogan QH, Stucky CL. Spinal Nerve Ligation in Mouse Upregulates TRPV1 Heat Function in Injured IB4-Positive Nociceptors. The Journal of Pain. 2010;11:588–99. [PubMed: 20015699]

51. Tingaker BK, EkmanOrdeberg G, Facer P, Irestedt L, Anand P. Influence of pregnancy and labor on the occurrence of nerve fibers expressing the capsaicin receptorTRPV1in human corpus and cervix uteri. Reprod Biol Endocrinol. 2008;6:8.

52. Yan T, Liu B, Du D, Eisenach JC, Tong C. Estrogen amplifies pain responses to uterine cervical distension in rats by altering transient receptor potential-1 function. Anesth Analg. 2007;104:1246–50.

53. Watanabe M, Ueda T, Shibata Y, Kumamoto N, Ugawa S. The role of TRPV1 channels in carrageenan-induced mechanical hyperalgesia in mice. Neuroreport. 2015;26:173–8.

54. Pierre MS, Reeh PW, Zimmermann K. Differential effects of TRPV channel block on polymodal activation of rat cutaneous nociceptors in vitro. Exp Brain Res. 2009;196:31–44.

55. Bautista DM, Movahed P, Hinman A, Axelsson HE, Sterner O, Högestätt ED, et al. Pungent products from garlic activate the sensory ion channel TRPA1, Proc Natl Acad Sci USA. 2005;102:12248–52.

56. Park CK, Kim K, Jung SJ, Kim MJ, Ahn DK, Hong SD, et al. Molecular mechanism for local anesthetic action of eugenol in the rat trigeminal system. Pain. 2009;144:84–94.

57. Ohkubo T, Shibata M. The selective capsaicin antagonist capsazepine abolishes the antinociceptive action of eugenol and guaiacol. J Dent Res. 1997;76:848–51.

58. Bandell M, Story GM, Hwang SW, Viswanath V, Eid SR, Petrus MJ, et al. Noxious cold ion channel TRPA1 is activated by pungent compounds and bradykinin. Neuron. 2004;41:849–57.

59. Peixoto-Neves D, Wang Q, Leal-Cardoso JH, Rossoni LV, Jaggar JH. Eugenol dilates mesenteric arteries and reduces systemic BP by activating endothelial cell TRPV4 channels. British Journal of Pharmacology. 2015; 172:3484–94

60. Choi TY, Park SY, Jo JY, et al. Endogenous expression of TRPV1 channel in cultured human melanocytes. J Dermatol Sci. 2009;56(2):128–30

61. Heiner I, Eisfeld J, Halaszovich CR, et al. Expression profile of the transient receptor potential (TRP) family in neutrophil granulocytes: Evidence for currents through long TRP channel 2 induced by ADP-ribose and NAD. Biochem J. 2003;371:1045–53.

62. Birder LA, Kanai AJ, de Groat WC, et al. Vanilloid receptor expression suggests a sensory role for urinary bladder epithelial cells. Proc Natl Acad Sci USA. 2001;98(23):13396–401.

63. Stander S, Moormann C, Schumacher M, et al. Expression of vanilloid receptor subtype 1 in cutaneous sensory nerve fibers, mast cells, and epithelial cells of appendage structures. Exp Dermatol. 2004;13(3):129–39.

64. Chen CW, Lee ST, Wu WT, Fu WM, Ho FM, Lin WW. Signal transduction for inhibition of inducible nitric oxide synthase and cyclooxygenase-2 induction by capsaicin and related analogs in macrophages. Br J Pharmacol. 2003;140(6):1077–87.

65. Mezey E, Toth ZE, Cortright DN, et al. Distribution of mRNA for vanilloid receptor subtype 1 (VR1), and VR1-like immunoreactivity, in the central nervous system of the rat and human. Proc Natl Acad Sci USA. 2000;97(7):3655–60.

66. Storozhuk, Maksim V., Moroz, Olesia F., and Zholos, Alexander V. Multifunctional TRPV1 Ion Channels in Physiology and Pathology with Focus on the Brain, Vasculature, and Some Visceral Systems. BioMed Research International. 2019;5806321.

67. Tominaga M, Caterina MJ, Malmberg AB, et al. The cloned capsaicin receptor integrates multiple pain-producing stimuli. Neuron. 1998;21(3):531–43.

68. Cromer BA, McIntyre P. Painful toxins acting at TRPV1. Toxicon. 2008;51(2):163–73.

69. Alawi K, Keeble J. The paradoxical role of the transient receptor poten-tial vanilloid 1 receptor in inflammation. Pharmacol Ther. 2010;125(2): 181–95.

70. Jung J, Hwang SW, Kwak J, et al. Capsaicin binds to the intracellular domain of the capsaicin-activated ion channel. J Neurosci. 1999;19(2):529–38.

71. Jordt SE, Tominaga M, Julius D. Acid potentiation of the capsaicin receptor determined by a key extracellular site. Proc Natl Acad Sci USA. 2000;97(14):8134–39.

72. Latorre R, Vargas G, Orta G, Brauchi S. Voltage and temperature gating in thermoTRP channels. In: Liedtke W, Heller S, editors. TRP Ion Channel Function in Sensory Transduction and Cellular Signaling Cascades. London, UK: CRC Taylor and Francis; 2007.

